# Diffuse fungal symbiosis in Deathwatch beetles

**DOI:** 10.64898/2026.08.10.743371

**Authors:** Austin Hendricks, T. Keith Philips, Tobias Engl, Rüdiger (Rudy) Plarre, Vincent G. Martinson

## Abstract

Many insects rely on symbiotic fungi to occupy specialized ecological niches, yet the evolutionary dynamics of these partnerships remain poorly resolved for most lineages. The beetle family Ptinidae, split into the morphologically distinct Spider beetles and Deathwatch beetles, has long been known to harbor fungal endosymbionts based on early microscopy, but few associations have been confirmed with molecular methods. Here, we combined ultra-conserved element (UCE) phylogenomics with ITS amplicon sequencing to test whether fungal endosymbionts are conserved across Ptinidae and whether they have cospeciated with their hosts. Our UCE phylogeny supports Spider beetles and Deathwatch beetles as monophyletic clades but indicates that some aspects of subfamily-level taxonomy may merit closer examination. Screening for three known symbiotic fungal genera (*Symbiotaphrina*, *Meyerozyma*, *Nakazawaea*) revealed *Symbiotaphrina* in most Deathwatch beetles but no Spider beetles, while the other two genera were present but uncommon. Despite widespread *Symbiotaphrina* infection, we found no phylogenetic mirroring between host and symbiont trees, indicating an absence of codiversification. Instead, distantly related hosts frequently shared closely related symbionts, consistent with diffuse, mixed-mode transmission involving both vertical and horizontal symbiont exchange. This pattern parallels those documented in fungus-farming termites, ambrosia beetles, ants, and woodwasps, suggesting that diffuse, mixed-mode symbiosis may be a general hallmark of long-term insect-fungal associations. We further identify an unidentified Helotiales group as a candidate novel endosymbiont, recovered consistently within a clade comprising *Anobium*, *Hemicoelus*, and *Ptilinus*. Together, these findings reframe Deathwatch beetle-fungal associations as a dynamic, evolutionarily labile symbiosis rather than a fixed partnership.

## Introduction

Insects are the most species-rich group of animals, a diversification driven in part by intimate associations with beneficial microorganisms that have enabled many lineages to exploit a broad array of ecological niches (Moran et al. 2005, Klepzig et al. 2009, Sudakaran et al. 2017). Within insects, obligate mutualistic microbes that supply essential nutrients have independently evolved more than 16 times, with bacteria comprising the vast majority of these associations (Cornwallis et al. 2023). These symbionts are typically transmitted vertically from parent to offspring, a process that preserves the association, aligns the fitness of symbiont and host, and results in codiversification (Bennett and Moran 2015). Among bacteria-insect symbioses, codiversification is widespread and occurs in associations involving both intracellular and extracellular symbionts (Salem et al. 2015). However, vertical transmission also subjects symbionts to repeated population bottlenecks that –in bacteria– frequently lead to reduced functional capacity (e.g., essential gene loss or pseudogenization) that, over evolutionary timescales, can result in the replacement of the symbiont by a new microbial partner (Bennett and Moran 2015, McCutcheon et al. 2019, Husnik and Keeling 2019). In contrast, fungal symbionts are primarily extracellular, with only a few known examples of intracellular mutualists restricted to a small number of insect lineages (Blackwell 2017, Matsuura et al. 2018). The evolutionary origins and dynamics of these fungal associations remain largely unresolved (Buchner 1965a, Blackwell 2017, Biedermann and Vega 2020).

Intracellular fungal symbionts can be divided loosely into two categories: 1) Ancestral mutualisms– host insects have no known obligate bacterial symbionts and have specialized organs adapted to house intracellular fungi (observed in flower longhorn beetles and Deathwatch beetles), and 2) Replacement mutualisms– a fungal pathogen of insects has supplanted an ancestral bacterial symbiont and are generally maintained outside of bacteriomes in host fat bodies or sheath cells (observed in some aphids, soft scales, planthoppers, leafhoppers, and cicadas) (Wang et al. 2022, Sasakura et al. 2024). Unlike bacterial symbionts, which often show phylogenies that mirror those of their insect hosts, intracellular fungal symbionts studied to date (cicadas and flower longhorn beetles) lack evidence of strict codiversification with their hosts (Wang et al. 2022, Sasakura et al. 2024). However, additional examples are needed to evaluate the generality of this pattern. One understudied group is the Deathwatch beetles.

Ancestral intracellular fungal mutualisms evolved independently in lineages of two distantly related beetle superfamilies—Flower Longhorn beetles (Chrysomeloidea; Cerambycidae; Lepturinae) and Deathwatch beetles (Bostrichoidea; Ptinidae *s.l.*; Anobiidae *s.s.*)—which have a common ancestor approximately 250 million years ago (McKenna et al. 2019). Despite being unrelated, these lineages exhibit strikingly convergent life-histories and physiological adaptations for maintaining fungal symbionts across both larval and adult stages: larvae possess anterior midgut mycetomes housing fungi within specialized cells called mycetocytes, while adult females have accessory organs that deposit extracellular fungal symbionts onto eggs during oviposition, ensuring vertical transmission to the next generation (Schomann 1937, Buchner 1965b, Kishigami et al. 2023). Notably, these ancestral fungal symbionts alternate between intracellular and extracellular phases within a single host generation—an uncommon life history for bacterial symbionts (Nardon and Grenier 1989). The extracellular phase of these fungal symbionts may profoundly influence the evolutionary dynamics of the symbiosis by increasing the effective population size during transmission bottlenecks and enabling hosts to acquire novel symbionts on ecological timescales via horizontal transmission.

Here, we focus on the beetle family Ptinidae *s.l*., which is generally split into two groups: the Spider beetles (Ptinidae *s.s.*) and the Deathwatch beetles (Anobiidae *s.s.*) (Table 1) (Philips 2002, Bell and Philips 2012a). Of the more than 2,200 described species, no Spider beetles have been screened for fungal symbionts, and only five species of Deathwatch beetles have been examined using DNA-based methods. Remarkably, these limited surveys have revealed three distinct genera of fungal symbionts: *Symbiotaphrina* (in *Stegobium paniceum* and *Lasioderma serricorne*), *Nakazawaea* (in *Ernobius mollis* and *Ernobius abietis*), and *Meyerozyma* (in *Xestobium plumbeum*) (Buchner 1921, Heitz 1927, Breitsprecher 1928, Müller 1934, Gräbner 1954, Jurzitza 1970). While *Symbiotaphrina* is a Pezizomycotina, the other symbionts— *Nakazawaea* and *Meyerozyma*—are members of the Saccharomycotina, with a most recent common ancestor dating to ∼520 Mya (Shen et al. 2020). This deep phylogenetic split strongly suggests that symbiont replacement has occurred in some Deathwatch beetles. The closely related family, Bostrichidae, is host to at least two ancestral bacterial endosymbionts (Engl et al. 2020, Kiefer et al. 2023); however, previous work found no evidence of conserved bacterial endosymbionts in either Spider beetles or Deathwatch beetles (Hendricks et al. 2025). Recently, a survey of *L. serricorne* and *St. paniceum* showed that both species are not restricted to one species of *Symbiotaphrina*, as different populations of each were host to different symbionts (Alina et al. 2026). Overall, it remains unknown whether Ptinidae harbors a single highly conserved fungal symbiont with strict codivergence, multiple lineage-specific symbionts, or diffuse associations with a fungal lineage punctuated by replacement events. Furthermore, the vast majority of Spider beetles and Deathwatch beetles remain un-surveyed, and a robust phylogeny for these lineages is lacking, limiting the ability to conduct cophylogenetic analyses.

**Table 1.**
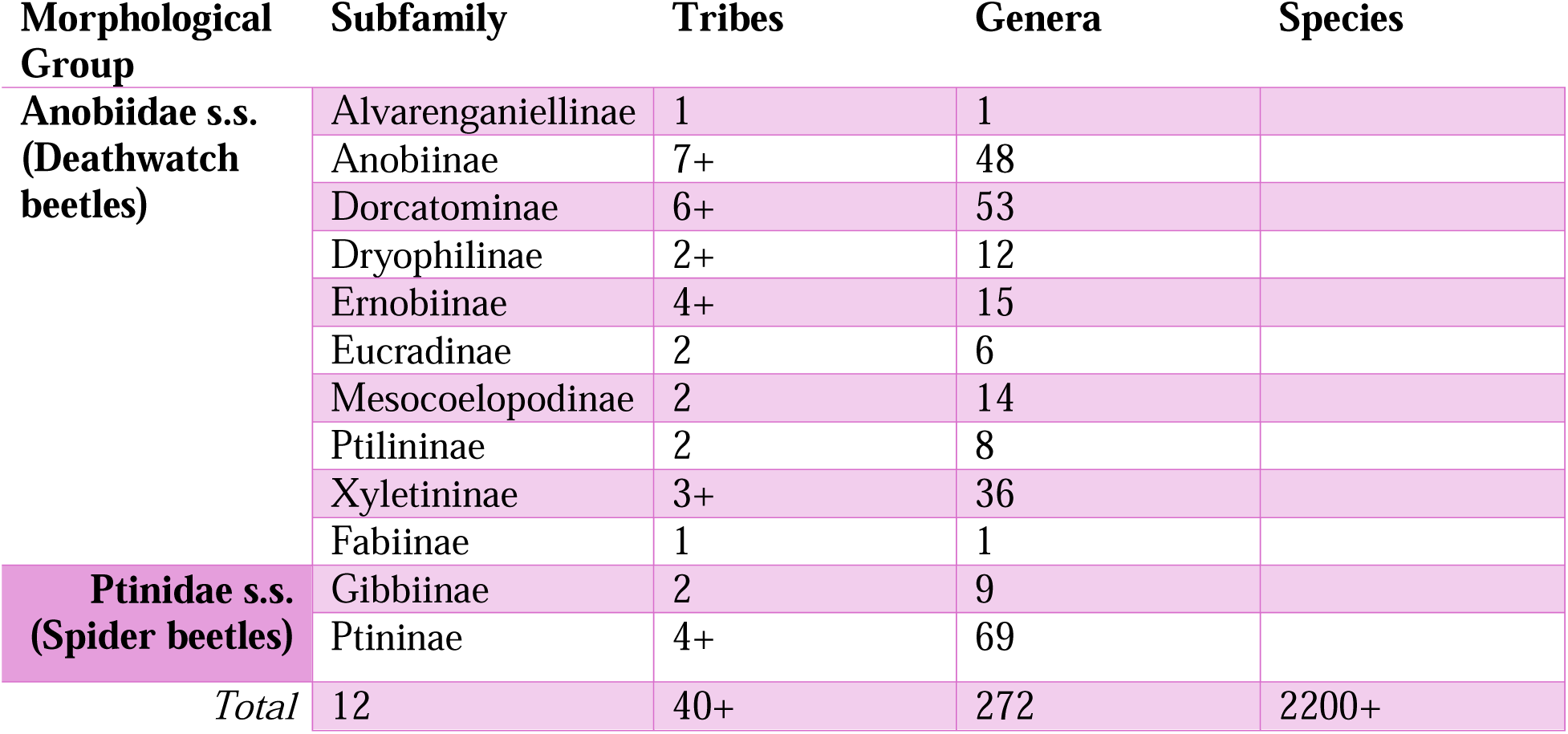
Taxonomic groups of Ptinidae.

In this study, we construct a phylogeny for Ptinidae based on genome-wide SNP data using ultra-conserved element (UCE) sequencing and describe the putative relationships with fungal endosymbionts through amplicon sequencing of the ITS ribosomal region. Our analyses center on the previously recognized symbiotic genera *Symbiotaphrina*, *Nakazawaea*, and *Meyerozyma*.

## Results

### UCE phylogeny for Ptinidae *s.l*

Earlier efforts to reconstruct the Ptinidae phylogeny left many relationships unresolved (Bell and Philips 2012b, Gearner 2019a); therefore, we began by constructing a genome-wide dataset using a Coleoptera-specific UCE probe set. Across the 43 specimens included in the final analyses, an average of 65% (761/1172) of the target UCE loci were captured. With this UCE dataset, we generated three concatenated matrices for phylogenetic analysis: a 50% matrix (50p), requiring each locus to be present in at least half of the specimens; a 75% matrix (75p); and a 95% matrix (95p). Although the 95p dataset had the most complete alignment, it contained few loci (48/1172), and phylogenetic analyses produced largely unresolved polytomies (Figure S1). Consequently, subsequent analyses focused on the 75p and 50p datasets to infer phylogenies using IQ-TREE2 (maximum likelihood, ML) (Figure S2), SVDQuartets (species tree from site patterns) (Figure S3), and ASTRAL (species tree from gene trees) (Figure S4). In all three analyses, the 75p and 50p datasets largely agreed with each other. For the remaining analysis, we chose to focus on the 50p dataset, which had the best overall bootstrap support values.

Nearly all trees supported the two historically recognized groups of Ptinidae, Deathwatch beetles and Spider beetles, as two reciprocally monophyletic clades. One exception, the 50p ASTRAL tree (Figure 1, S3), nested the Spider beetles within the Deathwatch beetles with Ernobiinae sister to all other Ptinidae (Figure 1, S1-S4). While it has been hypothesized that Spider beetles are a highly derived group of Deathwatch beetles (Crowson 1981, Ivie 1985), this arrangement is unlikely given that several relationships drop to extremely low bootstrap support (e.g., 3% for relationship between Ptinidae *s.s.* and Deathwatch beetles without Ernobiinae).

**Figure 1:**
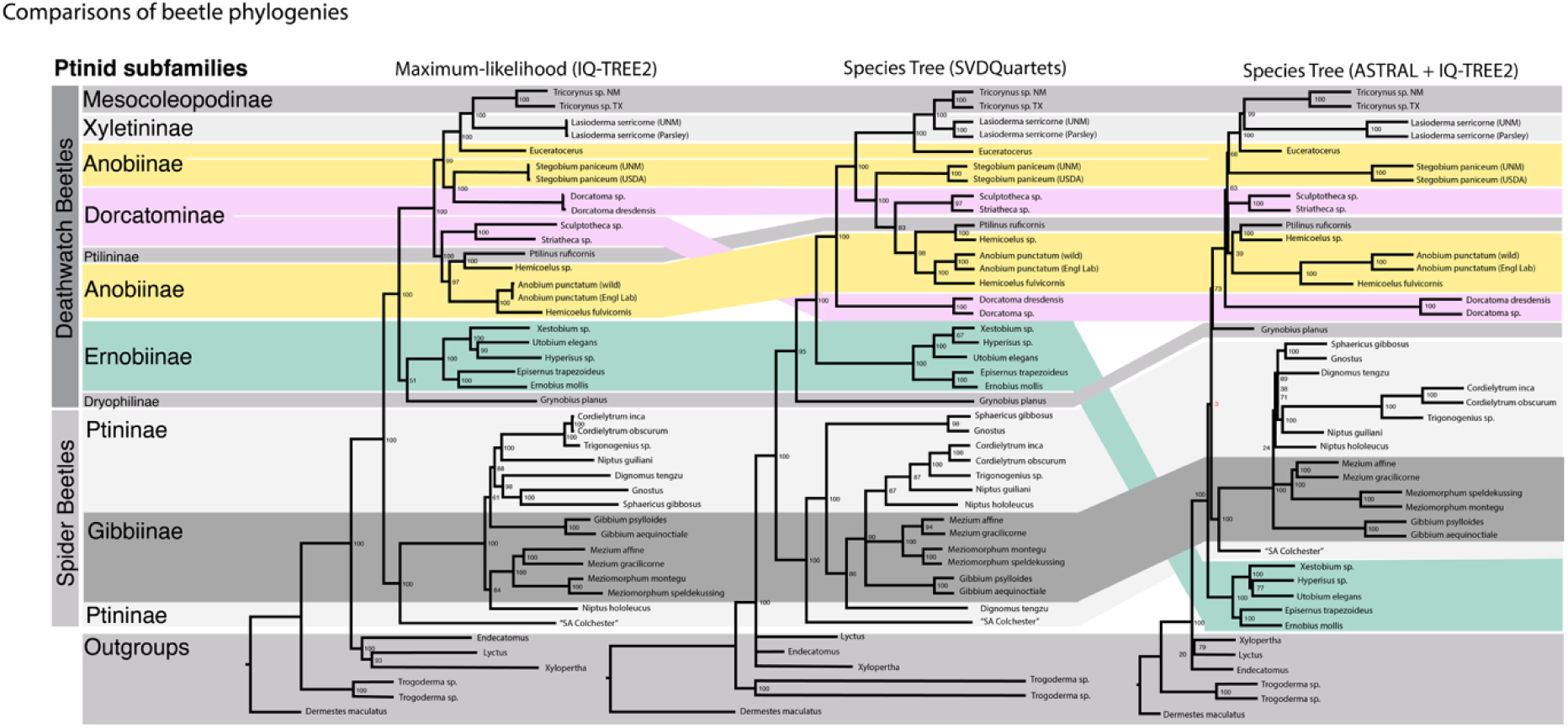
Phylogeny of beetle family Ptinidae. Comparison of three different methods: a maximum-likelihood tree created with IQ-TREE2, a species tree from site patterns with SVDQuartets, and a species tree from gene trees with ASTRAL. All trees are based on Coleoptera-specific UCE loci recovered from at least 50 percent of specimens (50p matrix). To better denote differences between methods, the subfamilies are color-coded. Support values for IQ-TREE2 are bootstraps, for SVDQuartets are bootstraps, and ASTRAL are concordance vectors.

Current taxonomic groupings were inconsistently supported by our UCE phylogenetic analyses. *Ernobiinae* was the only subfamily consistently recovered as monophyletic across all analyses; *Grynobius planus* (the sole representative of Dryophilinae) was generally placed as sister to Ernobiinae (Figures 1, S1–S4). Several groupings not recognized by current taxonomy were nonetheless consistently recovered as monophyletic, suggesting previously unrecognized close relationships: (1) *Tricorynus*, *Lasioderma*, and *Euceratocerus*; and (2) *Sculptotheca*, *Striatheca*, *Ptilinus*, *Hemicoelus*, and *Anobium*. In contrast, several subfamilies and genera were consistently split into two or more clades across analyses — including Dorcatominae, Anobiinae, and *Hemicoelus*. Overall, these results suggest that current taxonomic boundaries within Ptinidae may require revision, though increased taxon sampling will be necessary to confirm and refine these findings. Detailed phylogenetic results for both Deathwatch beetles and Spider beetles are provided in the Supplemental Results.

### Identification of known fungal associates

We used ITS amplicon sequencing to identify putative fungal symbionts, first focusing on sequences from the known symbiont genera *Symbiotaphrina*, *Nakazawaea*, and *Meyerozyma*.

Here, we will use ‘detected’ to indicate an ASV was found in a sample (at any amount), and significant for ASVs that made up over 1% of the relative abundance of a given sample. Not all samples produced an ITS library, so some specimens are missing fungal data. We detected *Symbiotaphrina* in 24/39 species screened (20/22 species of Deathwatch beetles, 5/15 species of Spider beetles, and 2/2 outgroups; Figure 2). Of those, 17/22 Deathwatch beetles, 2/15 Spider beetles, and 1/2 outgroups had significant portions of *Symbiotaphrina*. All specimens of the known hosts *L. serricorne* and *St. paniceum* had over 10% of ASVs corresponding to *Symbiotaphrina* (Figure 3). *Nakazawaea* and *Meyerozyma* were less common. *Nakazawaea* was detected in 13/39 species (9/22 species of Deathwatch beetles, 2/15 species of Spider beetles, and 0/2 outgroups; Figure 2) with only 4 species having it at significant abundance: 3 Deathwatch beetles (including its known host, *Ernobius mollis*) and 1 Spider beetle. Finally, *Meyerozyma* was detected in 10/39 species (9/22 species of Deathwatch beetles, 1/15 species of Spider beetles, and 0/2 outgroups; Figure 2). While *Meyerozyma* was not detected at high percentages in its canonical host *Xestobium* (0.07% of reads), it was found in the closely related *Hyperisus* (11.8% of reads) and in *Ptilinus ruficornis* (1.4% of reads) (Figure 2, 3).

**Figure 2.**
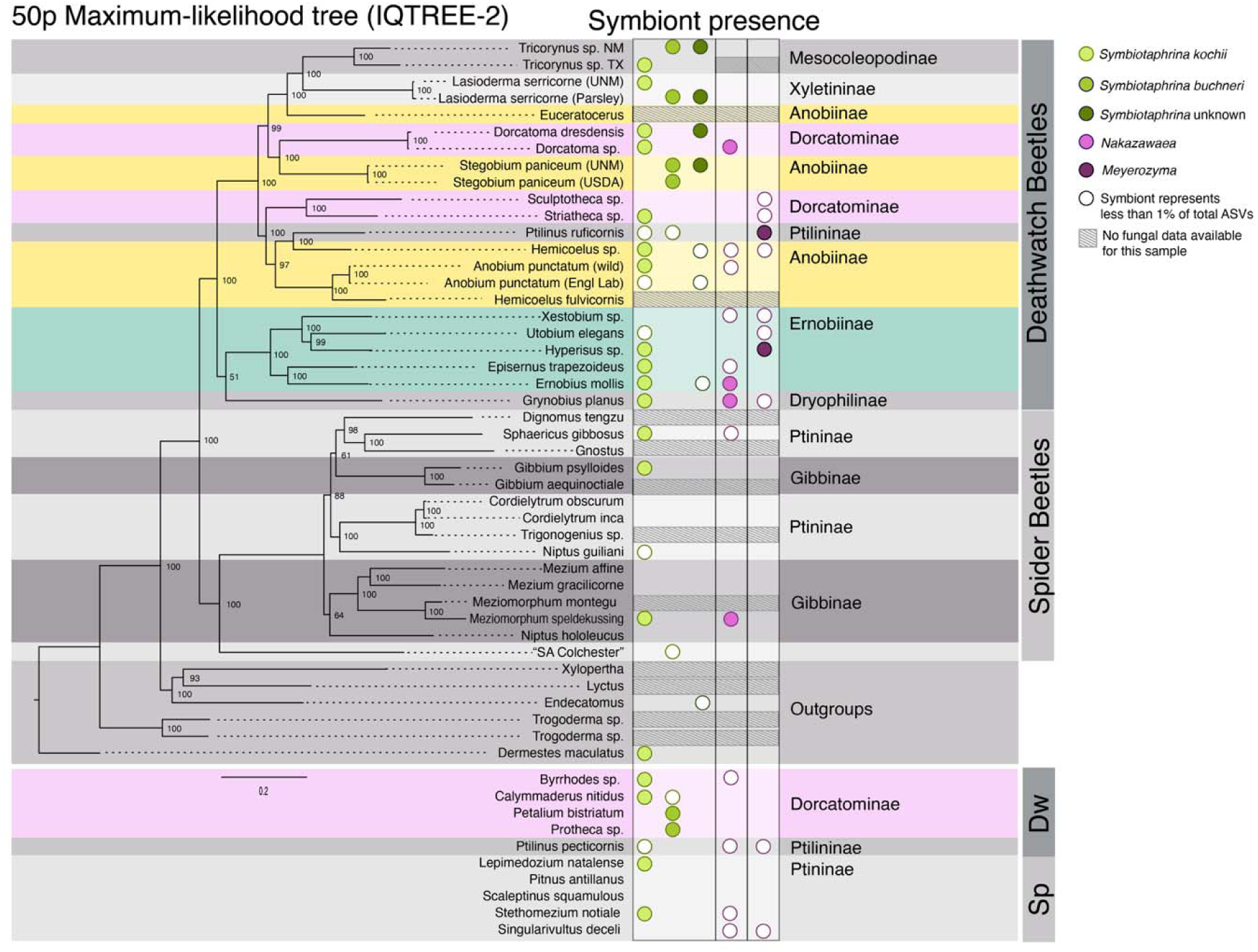
Fungal symbiont associations across the Ptinidae phylogeny. The 50p maximum-likelihood tree with symbiont presence indicated by a color-coded circle on the right. Symbionts identified from low-abundance ASVs (less than 1% of total ASVs for that sample) are colored white, and hosts with no ITS data are indicated with hashed lines.

**Figure 3:**
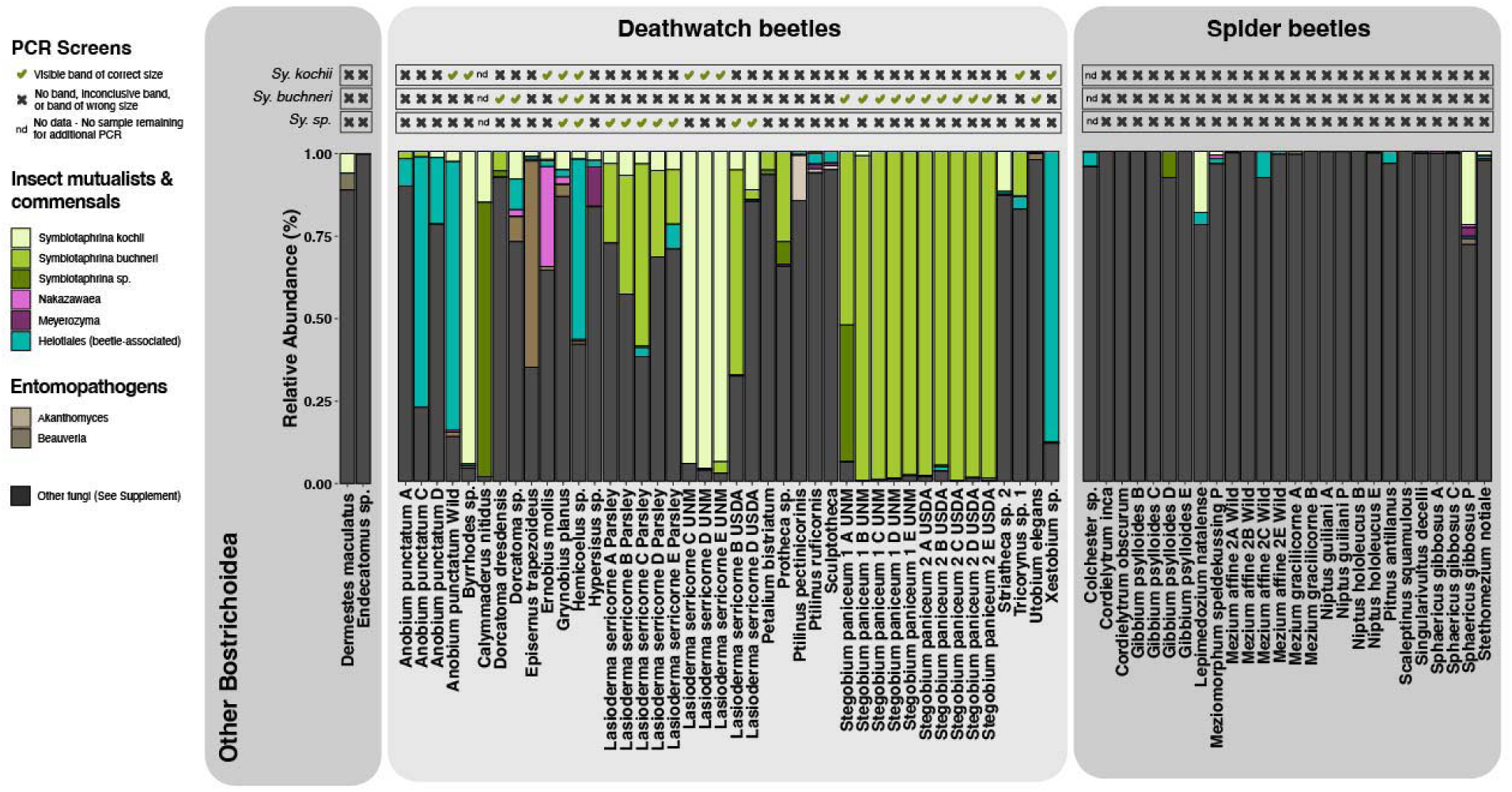
Relative abundances of insect-associated symbionts. Each column of the bar plot represents data from an individual beetle specimen. Insect-associated symbionts are color-coded. All remaining fungi are listed under “other”. The top three rows show the results of the PCR screen of each individual using *Symbiotaphrina* species specific primers. If the sample was not available for further screening, it was labeled ‘nd’ for no data.

### Phylogenetic analyses of fungal symbiont genera: *Symbiotaphrina*, *Nakazawaea*, *Meyerozyma*

For each fungal genus, we aligned all ASVs (ITS amplicon data) against representative GenBank sequences from that genus, encompassing both symbiotic and free-living (e.g., environmental or plant-associated) taxa. In total, 137 *Symbiotaphrina* ASVs, 12 *Nakazawaea* ASVs, and 15 *Meyerozyma* ASVs were identified by Qiime2, confirmed by BLAST to the nr database, and used to create phylogenetic trees (Figure 4, S5). Relationships among *Symbiotaphrina* species differed somewhat from a previous phylogeny based on full-length SSU, ITS, and LSU sequences (Baral et al. 2018), which likely reflects the shorter ITS fragments used in our alignment (227–541 bp). Two ASVs were initially identified as the plant-associated species *Sy. lignicola*; however, they had very low read counts (less than 0.016% of reads in the sample) and they may have been the result of sequencing error, contamination, or environmental exposure. All other *Symbiotaphrina* ASVs from our study clustered into three groups corresponding to known Deathwatch beetle-symbiont species: *Sy. kochii*, *Sy. buchneri*, and a novel *Symbiotaphrina* sp. first identified in Alina et al. (2026) (Figure 4). Similar to the pattern observed in *Symbiotaphrina*, all 12 *Nakazawaea* ASVs formed a clade with *Nakazawaea ernobii* sequences, a previously known Deathwatch beetle-symbiont (Figure S5a); and all 15 *Meyerozyma* ASVs formed a clade corresponding to the *M. carpophila* and *M. xestobii* species complex, which includes isolates from Deathwatch beetles (Romi et al. 2014) (Figure S5b).

**Figure 4:**
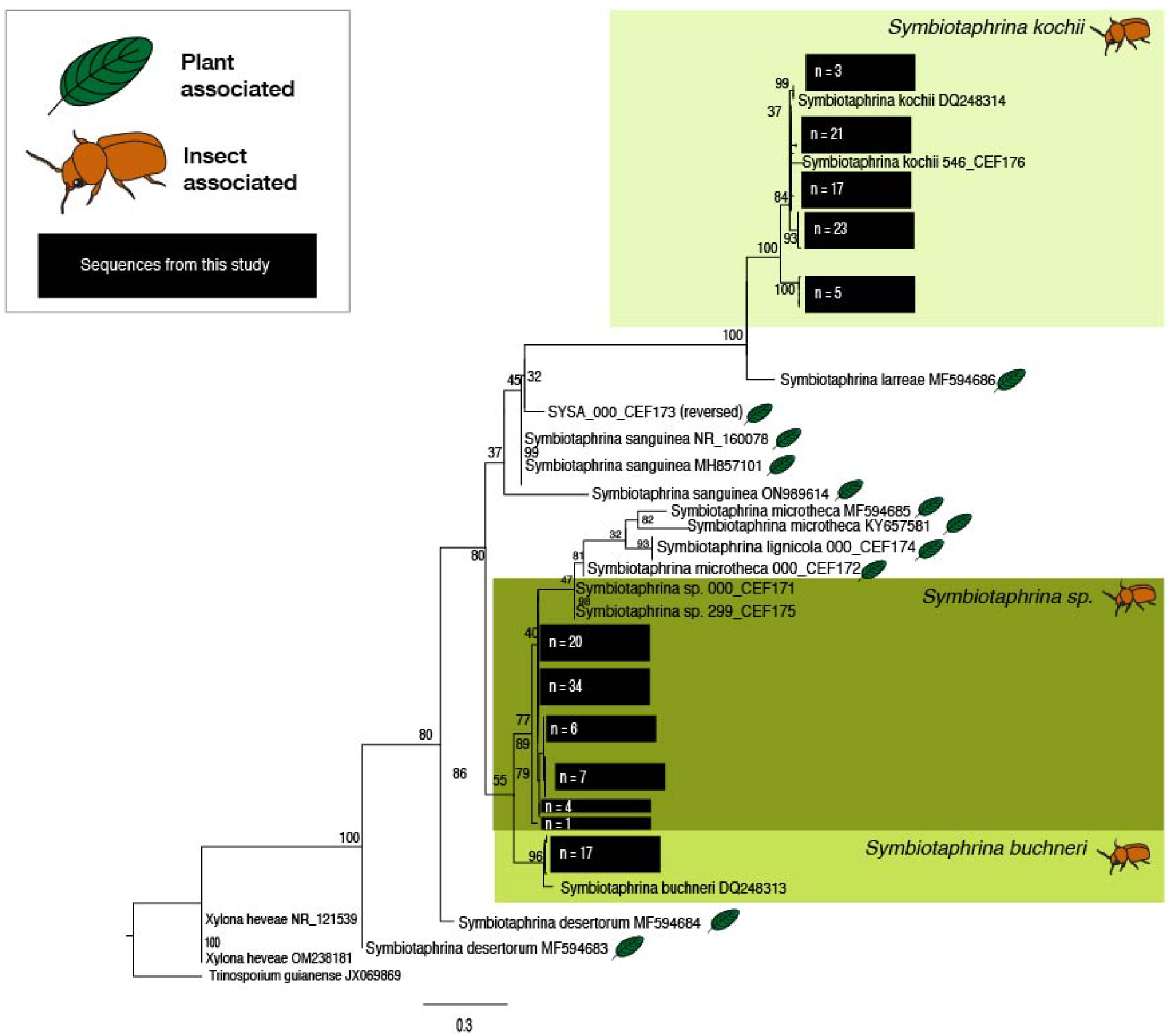
Phylogeny of the fungal genus *Symbiotaphrina* including ASVs recovered from beetle survey. The maximum-likelihood tree (IQ-TREE2) is based on short amplicon ITS sequences and has bootstrap support values. Representative ITS sequences from each *Symbiotaphrina* species are labeled and sequences from the beetle survey are in black boxes. Species are denoted as being either plant-associated or insect-associated.

### Verification of *Symbiotaphrina* species

While ITS amplicon sequencing offers an unbiased window into the fungal associations of beetles, it has several limitations: short sequences that obscure closely related species, potential for contamination, and swamping of samples by highly abundant fungi — all of which can complicate the scoring of symbiont presence and absence. In particular, our ITS data suggested that some specimens may be harboring multiple species of *Symbiotaphrina* (Figure 3), which has not been previously confirmed. To verify these associations, we designed species-specific PCR primers to differentiate *Symbiotaphrina* species (Figures 3, S6). Although DNA from two specimens was exhausted prior to screening, the rest were successfully screened, providing a clear picture of *Symbiotaphrina* associations.

ITS amplicon screening detected *Sy. kochii* in 22/39 species screened (16/22 species of Deathwatch beetles, 6/15 species of Spider beetles, and ½ outgroups; Figure 2, 3). However, PCR screening using the *Sy. kochii* primers only confirmed *Sy. kochii* in 8 Deathwatch beetle species (Figure 3). A similar pattern followed for *Sy. buchneri -* it was detected by ITS screening in 7/39 species (6/22 species of Deathwatch beetles, 1/15 species of Spider beetles, and 0/2 outgroups) – but was only confirmed in 6 Deathwatch beetle species. Finally, the novel *Symbiotaphrina* was not identified by Qiime2, likely due to the lack of representation in its database. Unidentified *Symbiotaphrina* ASVs were found in 13/39 species (11/20 species of Deathwatch beetles, 1/20 species of Spider beetles, and 1 /2 outgroups), but only 3 Deathwatch beetle species tested positive for the novel *Symbiotaphrina* with PCR primers. Within the ITS dataset, the novel *Symbiotaphrina* appears to have been misidentified as *Sy. kochii* or *Sy. buchneri* in some samples, again, likely connected to lack of available data in the Silva dataset used. Overall, our PCR assays confirmed the presence of *Symbiotaphrina* only in Deathwatch beetles, whereas it was not detected in any Spider beetles.

Of the species screened, only *Grynobius planus* and *Hemicoelus* sp. showed evidence of co-infection, testing positive across all three *Symbiotaphrina* primer sets (Figures 3, S6); however, ITS analysis of both specimens revealed that *Symbiotaphrina* reads corresponded predominantly to *Sy. kochii* (Figure 2, 3). Although the primers were validated against numerous plant-associated *Symbiotaphrina* species to rule out cross-reactivity (S6), off-target amplification of non-*Symbiotaphrina* taxa may explain these results, particularly for the *Sy. buchneri* and *Sy.* ‘novel’ primer sets. While co-infection by multiple *Symbiotaphrina* species cannot be fully ruled out in these specimens, further work is needed to verify these results.

### Identification of a potential novel fungal symbiont

Following analysis of the known symbiont genera, we evaluated the remaining fungal reads for evidence of additional candidate symbionts, restricting our analysis to fungal genera with known insect associations that 1) were detected in more than one species in our dataset and 2) comprised ≥5% of the fungal reads in at least one specimen. We identified two members of Cordycipitaceae known to cause pathogenic infections in beetles: *Beauveria* (*Dermestes maculutaus, Dorcatoma sp*., *Episernus trapezoideus*) and *Akanthomyces* (*Ptilinus pectinicorinis, Ptilinus ruficornis*, *Sculptotheca sp.*) (Figure 3) (Imoulan et al. 2017, Wang et al. 2024). We also found a group of 56 unidentified Helotiales (Leotiomycetes) ASVs that had a closest match (90% identity) in GenBank to OP537775.1, an unclassified fungal isolate from the oviduct of a Flower Longhorn beetle (Sasakura et al. 2022). On alignment and clustering, this group formed 3 clusters of sequences with nearly 100% identity (Figure S7). From an ITS tree that included other Leotiomycetes species (Figure S7), OP537775 clustered within our group of sequences. These Helotiales ASVs were abundant in all examined specimens of *Anobium punctatum* (4/4), including both wild and lab-reared individuals, as well as in *Hemicoelus* and *Xestobium*. This Helotiales ASV also occurred at lower abundance in 8 additional Deathwatch beetle species and 5 Spider beetle species (Figure 3). More work is required to fully identify this Helotiales taxon and evaluate its status as a symbiont.

Other common fungal families in the dataset included ASVs identified from Cladosporiaceae, Tremellaceae, Hyaloscyphacaea, Aspergillaceae, Dothioraceae, and Saccharomycetales (Figure S8), all of which are common in the environment and encompass many lifestyles. After the genera identified in Figure 3, the most common genera were *Cladosporium*, *Cryptococcus, Aspergillus, Dothiora,* and *Penicillium* (Supplementary Data).

### Cophylogeny test for *Symbiotaphrina*

Because *Symbiotaphrina* symbionts are known to be vertically transmitted and are abundant in this dataset, we also attempted a global-fit test in PaCO as a measure of codiversification. The result was an m^2^ of 27.4 with 1000 permutations, indicating no congruence between the host tree (50p ML beetle phylogeny, Figure 1A) and the symbiont tree (Figure 4).

## Discussion

In this study, we set out to map the relationships between Ptinidae beetles and their fungal symbionts. To accomplish this, we constructed a new host phylogeny using ultra-conserved element sequencing on 43 specimens, screened 70 specimens for fungal associates using high-throughput amplicon sequencing, and confirmed the presence of three *Symbiotaphrina* species through specific primer sets. We found that in most analyses, the Deathwatch beetles and Spider beetles form two monophyletic clades, and that many subfamilies may need to be revisited with molecular data in the future. From ITS screening and confirmation PCR, we found that nearly all Deathwatch beetles screened had evidence for one or more known fungal symbionts, and that while ITS screening detected the presence of some symbionts in Spider beetles, none were confirmed via PCR. All high abundance sequences for the three symbiotic genera *Symbiotaphrina, Nakazawaea,* and *Meyerozyma* clustered with insect-associated species, and not the free-living relatives that are present in the environment. Therefore, we conclude that most, if not all, Deathwatch beetles harbor a fungal endosymbiont; however, unlike bacterial endosymbionts, there does not appear to be a one-to-one, conserved host-endosymbiont relationship, but rather a diffuse association with several taxonomically restricted fungal taxa.

### Spider beetles and Deathwatch beetles form reciprocally monophyletic clades

There are striking differences between Deathwatch beetles and Spider beetle in their external morphology, dietary preferences, and interaction with fungal symbionts. Spider beetles are named for their round abdomen and long legs, which give them a superficial resemblance to spiders. They are frequently scavengers or dung-feeders, a diet that fulfills their nutritional requirements (Andrews 1967, Aalbu and Andrews 1992). Additionally, no internal structures associated with maintaining a fungal symbiont (i.e., mycetome, accessory organs) have been observed in this group. Deathwatch beetles have more cylindrical body plans with the head bent downward under a cowl-like pronotum, more reminiscent of other wood-boring beetle taxa. Many taxa subsist on dry wood or dry shelf fungi, which provide low nutrition and likely require additional nutrients from a symbiotic microbe to survive (similar to many other wood-boring taxa) (Cornwallis et al. 2023). All Deathwatch beetles dissected have been found to have mycetome and accessory structures that are used to maintain a fungal mutualist (intracellular in the mycetome) (Martinson 2020). Further supporting monophyly for these two beetle groups is the distribution of mutualistic fungi we found through ITS surveys and PCR screening. Most Deathwatch beetles screened had ITS sequences corresponding to mutualistic fungi, and PCR screening confirmed *Symbiotaphrina* in many of these species. In striking contrast, *Symbiotaphrina* was not detected in any Spider beetles by PCR screening.

Morphology-based analyses reconstruct reciprocally monophyletic clades (Li et al. 2023); yet, single and multi-gene trees were previously unable to resolve this relationship (Gearner 2019b). Our UCE analyses produced trees supporting reciprocal monophyly in nearly all our analyses (Figure 1, S1-S4), and with as few as 48 loci (95p matrix) we found strong support (Figure S1). The one exception, the 50p ASTRAL tree, placed Ernobiinae sister to all other Ptinidae, but with extremely low bootstrap support (3%) – indicating that method had considerable uncertainty in the outcome.

Altogether, it remains possible that Spider beetles are highly divergent members of the Anobiidae (Crowson 1981, Ivie 1985); however, the plurality of morphological, symbiotic, and sequence data instead indicate that fungal symbiosis evolved independently within Deathwatch beetles, and that Spider beetles form a separate monophyletic group. Resolving the relationships, and potentially the taxonomic classifications, within these groups will require additional work, particularly greater taxon sampling, given that 1) some subfamilies (i.e., Xyletininae, Dryophilinae) are represented by only a single species, and 2) other subfamilies (e.g., Anobiinae, Dorcatominae) are consistently split across all analyses (Figures 1, S1–S4).

### *Symbiotaphrina* associations differ by population

While *L. serricorne* and *St. paniceum* were already known to harbor *Symbiotaphrina*, we show that this symbiont is broadly associated with diverse Deathwatch beetles (Figure 2, 3). Our findings also support those from Alina et al. (2026), which found that different populations of *L. serricorne* and *St. paniceum* can harbor different symbionts. While Alina et al. (2026) found *L. serricorne* associated with *Sy. kochii* or *Sy. buchneri* and *St. paniceum* associated with the novel *Symbiotaphrina*, we found *L. serricorne* associated with *Sy. kochii* or the novel *Symbiotaphrina*, and *St. paniceum* associated with *Sy. buchneri* (Figure 3). Additionally, *Ernobius* was associated with *Sy. kochii* in our survey, but the novel *Symbiotaphrina* in Alina et al. (2026). Furthermore, symbiont exchange between *L. serricorne* and *St. paniceum* has been experimentally validated; Pant and Fraenkel (1954) removed the native symbiont by surface sterilizing the egg and subsequently feeding the aposymbiotic larvae the alternative symbiont. Together, these observations indicate that symbiont replacement occurs relatively frequently, on ecological rather than evolutionary time scales.

Given these symbiont replacements, one might expect co-infections to occur; however, our data suggest that co-infection by multiple *Symbiotaphrina* species is rare. Our PCR survey found only two specimens with possible co-infections, *Grynobius planus* and *Hemicoelus sp.*, but we lacked multiple individuals to screen and we could not rule out the possibility of primer cross-reactivity to non-*Symbiotaphrina* fungi. If co-infections *were* common, we would expect to find them in wild pest populations, which seem to have frequent enough crossover to switch symbionts entirely. However, across all 8 of the populations of *L. serricorne* and *St. paniceum* surveyed – both the current study and in Alina et al. (2026) – individuals were found to maintain only one symbiont at a time. This pattern may result from 1) priority effects, whereby the initial symbiont to colonize a host excludes subsequent colonization, or 2) competitive exclusion between symbionts.

### Evolutionary implications of *Nakazawaea* and *Meyerozyma* infections

While previously described as primary symbionts and cultivated from Deathwatch beetles, *Nakazawaea* and *Meyerozyma* were found in high abundance in very few species (4 and 2, respectively). Furthermore, ITS amplicon data and PCR screening for *Symbiotaphrina* showed that these two fungal genera were commonly co-infections with other symbionts/potential symbionts: *Ernobius mollis* was co-infected with *Sy. kochii* and *Nakazawaea*, *Ptilinus ruficornis* was co-infected with *Akanthomyces* and *Meyerozyma, Hyperisus* was co-infected with *Sy. kochii* (no PCR confirmation due to lack of DNA) and *Meyerozyma* (Figure 3). Increased taxon sampling across all Deathwatch beetles, and increased number of specimens per species, would help map the host-range distribution of all symbionts and confirm co-infections by multiple endosymbiont genera. However, if *Nakazawaea* and *Meyerozyma* are genuinely less common, this may reflect greater host restriction—that is, an association limited to certain beetle species— or a more facultative symbiotic role, in which they provide benefits under some conditions but incur costs under others, leading to their loss.

### Identifying novel candidate fungal symbionts

Of the remaining fungal genera identified, few appear to be strong candidates for mutualistic symbiosis. *Aspergillus* and *Penicillium* species have been isolated from beetles, including *Lasioderma serricorne* (Kawakami and Takahashi 2007, Yoshinami et al. 2018), but there is currently no evidence of intracellular infection of the mycetome or vertical transmission for either genus. We included the Cordycipitaceae genera *Beauveria* and *Akanthomyces* in our analysis because members of these genera were recently found to provide defensive benefits to Dinidorid stink bugs, with fungal hyphae physically protecting eggs from parasitoid wasps (Nishino et al. 2025), and some *Ophiocordyceps* species have transitioned from parasites to mutualists in multiple insect lineages (Matsuura et al. 2018). Unfortunately, ITS sequences alone cannot resolve the type of relationship these Cordycipitaceae taxa have with their hosts; however, the absence of a consistent pattern of association among Deathwatch beetles, suggests a pathogenic rather than mutualistic interaction is more likely. The most promising novel symbiont candidate is an unidentified Helotiales group, as many of these sequences were recovered from the same host clade, comprising *Anobium, Hemicoelus,* and *Ptilinus* (Figure 2, 3); this pattern is consistent with vertical transmission. Targeted PCR amplification and sequencing of longer gene regions, examination of live specimens, and culturing will be required to further investigate this group and confirm these findings.

### Diffuse symbiosis as a hallmark of insect-fungal associations

Across the Deathwatch beetles surveyed, we observe a pattern in which distantly related hosts shared closely related symbionts. This is most evident in the widespread occurrence of *Sy. kochii* across all major lineages of our UCE phylogeny. This pattern is not consistent with environmental acquisition, as the fungal symbionts recovered were not simply any fungus, but instead fell into discrete species already known to be symbionts; this was true across *Symbiotaphrina*, *Nakazawaea*, and *Meyerozyma* (Figure 4, S5, S6). These findings are consistent with the concept of ‘diffuse symbiosis’, in which multiple host species share closely related symbiont taxa that form a pool of interchangeable genotypes capable of associating with diverse hosts, rather than showing the one-to-one codiversification expected under strict vertical transmission. This pattern may be best explained by mixed-mode transmission, in which both vertical and horizontal transmission occur and jointly shape the association between hosts and symbionts (Ebert 2013). Moreover, these symbiont taxa are rarely found in free-living conditions and instead appear restricted to host association, suggesting that they have become adapted, or effectively domesticated, to conditions provided by their hosts.

This pattern is paralleled in diverse insect systems with *extracellular* fungal symbionts. In fungus-farming termites, *Termitomyces* fungal cultivars form a monophyletic clade, indicating that the association is ancient and singular; yet individual termite lineages frequently switch between fungal partners within that clade rather than cospeciating with them (Aanen et al. 2002, Van De Peppel et al. 2021). Similar patterns have been documented in other fungus-farming insects, including ambrosia beetles (Mayers et al. 2020), ants (Schultz et al. 2024), and woodwasps (Hajek et al. 2013).

This general pattern extends to the more intimate, intracellular fungal symbioses found in some insects. Ancestral intracellular fungal symbionts—such as those of flower longhorn beetles and Deathwatch beetles—are transmitted to the next generation on the surface of the egg, a life history that creates an opportunity for horizontal transmission and symbiont replacement (Martinson 2020, Sasakura et al. 2024, Alina et al. 2026). By contrast, no extracellular stage is known for intracellular fungal symbionts that have replaced ancestral bacterial symbionts (e.g., *Ophiocordyceps*, which has replaced bacterial symbionts in diverse lineages). Nevertheless, a pattern of repeated symbiont replacement – involving multiple *Ophiocordyceps* lineages – is observed in cicadas, the one lineage with sufficient sampling to reveal this pattern (Wang et al. 2022). While additional surveys are needed, these patterns collectively suggest that frequent symbiont switching and replacement—generating diffuse symbioses—may be a hallmark of long-term insect-fungal associations.

## Materials and Methods

### Beetle specimens & UCE preparation, sequencing, and analysis

Insect specimens were obtained as previously published (Hendricks et al. 2025). Detailed information about specimen sources and DNA extraction can be found in Supplemental Table 1 (https://doi.org/10.6084/m9.figshare.33067949). For each beetle specimen, a shotgun library was constructed using the NEBNext Ultra II FS DNA Library Prep Kit for Illumina (Catalog # E78005, New England Biosciences, Ipswich, MA, USA). Samples were diluted for a total of 100 ng of input and incubated at 37°C for 8 minutes to achieve insert sizes between 300 – 700 bp. Adapters were not diluted prior to adapter ligation. Samples received 8-10 cycles of PCR (see Supplemental Table 1 for sample-specific information), and indexing was done using the NEB Unique Dual Index Primer Pairs (Catalog # E6440S). Samples were cleaned according to the standard protocol with the included beads. A subset of libraries was checked for sizing on an Agilent Bioanalyzer 2100 (Agilent Technologies, Santa Clara, CA), and library concentration was measured with the Qubit dsDNA HS kit (Catalog # Q32851, Thermofisher Scientific, Waltham, MA, USA). Most libraries (47/48) moved forward to UCE capture with the myBaits Coleoptera 1.1Kv1 bait set (Catalog # 305908.v5, Arbor Biosciences, Ann Arbor, MI, USA), which targets 1172 UCE loci (Faircloth 2017). Libraries were grouped based on concentration and pooled in sets of 7 or 8, with concentrations adjusted to meet the recommended input of ∼2 µg per pool. Each pool was concentrated down to 7 µl using NEBNext Sample Purification Beads (Catalog #E6178S). The initial hybridization was done at 62 °C for 24 hours. From there, we followed the manufacturer’s recommendations for UCE capture. Enriched libraries were amplified using the NEB Ultra II Master Mix for 10 cycles. Final evaluation was performed with the Bioanalyzer and Qubit, and enriched libraries were pooled equimolar and sequenced with a 300 bp paired-end P1 flow cell run on a NextSeq 2000 (Illumina, San Diego, CA).

Initial data processing was done using the PHYLUCE package v1.7.3 (Faircloth 2016). Briefly, default parameters were used to: clean raw reads were with Illumiprocessor (Bolger et al. 2014, Faircloth n.d.), assemble contigs with SPAdes (Bankevich et al. 2012), extract contigs matching UCEs with MAFFT (Katoh and Standley 2013), and trim resulting UCE contigs with GBLOCKS (Talavera and Castresana 2007, Castanera et al. 2017). Specimens yielded an average of 2.3 million reads (180,056–5,184,675), which assembled into about 34,000 contigs per sample (4,349–118,522) with a mean length of 318 bp (21–1,780 bp). From these assemblies, an average of 626 UCE loci were recovered per specimen (221–759). Basic statistics for the UCE matrices were calculated with a custom R script, that, briefly, takes listed loci from Nexus files generated by phyluce and creates a presence/absence matrix for each sample, in which a 1 represents the presence of the locus (but no quality statistics for any particular locus) and 0 represents the absence of a locus (<u>Figshare link to script</u>, Figure S9). Despite sufficient read coverage, four samples (*Bostrichus capucinus*, *Byrrhodes* sp., *Oligomeres ptilinoides*, and *Ptilinus pecticornis*) were excluded from further analysis due to low UCE locus recovery (Figure S9). Across all specimens, UCE enrichment averaged 2.8% of reads aligning to UCE loci (0.35– 13.57%); however, the four excluded samples were clear outliers, with substantially lower proportions of reads mapping to UCE loci, indicating they had poor probe enrichment during library preparation. For downstream phylogenetic analyses, three concatenated matrices were built using different parameters: a 50% matrix (50p), where at least half of the specimens were represented in each locus; a 75% matrix (75p); and a 95% matrix (95p). The 50p matrix included 763/1172 UCE loci (65%) with an alignment length of 179043 bp; the 75p matrix had 397 loci (34%) with 90756 bp; and the 95p matrix had 48 loci (4%) with 11159 bp. Partitioning schemes and substitution models for all three matrices were determined with sliding-window site characteristics (SWSC) in PartitionFinder2 (Lanfear et al. 2017, Tagliacollo and Lanfear 2018), resulting in 569, 350, and 70 partitions for the 50p, 75p, and 95p matrices, respectively. Maximum-likelihood (ML) trees were made using IQ-TREE2 with 1000 ultrafast bootstrap replicates (Minh et al. 2020). Bayesian inference (BI) analysis was made for the 95p matrix in MrBayes with 120,000 generations, discarding the initial 25% as burn-in (Ronquist and Huelsenbeck 2003). Multispecies coalescent phylogenies were built using default settings through the PAUP implementation of SVDQuartets (Chifman and Kubatko 2014, 2015) which infers species trees directly from site patterns using a quartet-based coalescent approach, and with ASTRAL (v.5.7.8) (Zhang et al. 2018) which reconstructs species trees from a set of gene trees by identifying the topology that maximizes shared quartet frequencies. SVDQuartets analysis evaluated 100,000 quartets, and 1000 bootstrap replicates were performed. For input into ASTRAL, individual gene trees were built in IQ-TREE2 using the MAFFT aligned, GBLOCKS trimmed alignments for each UCE locus. To assess support for the coalescent-based tree topology, we calculated gene concordance factors (gCF) and site concordance factors (sCF) in IQ-TREE2 and mapped the resulting quartet concordance vectors onto the ASTRAL phylogeny. The final concordance vectors were generated in R using Lanfear’s concordance_vector R script (Lanfear and Hahn 2024). Two groups of taxa showed positional instability consistent with long-branch attraction (LBA) in the ML trees: *Stegobium paniceum* and the *Dorcatoma* species. To test whether these taxa were artifactually displacing, we constructed two additional concatenated matrices using a 50% locus occupancy threshold; one excluding *St. paniceum* and one excluding *Dorcatoma* spp. Each matrix was analyzed in IQ-TREE2 under the same settings described above, without partitioning.

### Fungal ITS region sequencing and analysis

Library preparation was done at Novogene (Beijing, China) using the primer set ITS5-1737F and ITS2-2043R to target the ITS1 region of fungi. A total of 127 individual adult beetles were screened for possible symbionts. Of these, 70 individuals passed quality control and were moved forward to library preparation, covering 38 species of beetle. Initial analysis was done using the Qiime2 pipeline (2022.8) (Bolyen et al. 2019). The raw data was denoised and amplicon sequence variants (ASVs) were generated using Qiime2’s implementation of DADA2 with paired-end settings and a truncation length of 222 bp for forward reads and 224 bp for reverse reads (Bolyen et al. 2019). ASVs were classified using the UNITE database (10/05/2021 version) (Abarenkov et al. 2024). ASVs identified as fungi or Ascomycota with no lower taxonomic classifications were separated into a new file and run through BLAST to the NCBI nr database to ensure no ASVs were misclassified. ASVs were removed from further analysis if their top BLAST hit aligned to plants, animals, bacteria, or had a query cover under 20% - the last criteria covered a number of reads that otherwise were identified only as “Fungi” by Qiime2, and may be the result of inaccurate reads or chimerism. All sequences that were removed from the dataset are available at our Figshare (https://doi.org/10.6084/m9.figshare.33067949). To verify the taxonomic classification of sequences assigned to fungal genera that include known Deathwatch beetle symbionts – *Symbiotaphrina*, *Nakazawaea*, and *Meyerozyma* – sequences were independently run through BLAST to the nr database and then used to create genus-specific phylogenies. Briefly, sequences matching the focal symbiont genera were aligned with MUSCLE 3.8.425 (Edgar 2004) and a Neighbor Joining phylogeny (Jukes-Cantor model) with 1000 bootstrap replicates was created in Geneious Prime 2026.1.

### Specific primer screens for *Symbiotaphrina*

Because this fungal ITS dataset was derived from whole-body DNA extractions and symbiont reads often represented only a small proportion of total reads, we sought to confirm *Symbiotaphrina* species identifications through targeted PCR amplification with taxon-specific primers. We first tested the primers from Alina et al. (2026); however, they could not differentiate the plant-associated *Symbiotaphrina* species (*Sy. microtheca*, *Sy. lignicola*, *Sy. sanguinea*) from *Sy. buchneri* and the “novel” *Sy.* species. Therefore, we designed specific primer pairs capable of distinguishing each of the three symbiotic *Symbiotaphrina* species: *Sy. kochii, Sy. buchneri,* and the “novel” *Sy.* species. Primers were designed based on variable regions in the mitochondrial genomes of the three clades. Specificity of these primers was first confirmed *in silico* by using Geneious Prime 2026.1 to check the primers against all mitochondrial genomes in our dataset, allowing for up to three mismatches. To confirm specificity, candidate primers were ordered (IDT) and directly screened against a panel of fungal DNA isolates (i.e., *Sy. kochii*, *Sy. buchneri*, *Sy. sp ‘novel*’, *Sy. lignicola, Sy. microtheca, Sy. sanguinea*), as well as, an artificial mix of the three symbiotic species (Figure S6). Briefly, PCRs were done using NEB Taq (#M0273) and the manufacturer’s standard PCR mix recommendations. Cycling conditions were initial denaturation of 94°C for 2 min, then 35 cycles of 94°C for 20 s, variable annealing temperature for 20 s, 65°C for 50 s, and then a final extension of 65°C for 5 minutes, followed by a hold at 4°C. To confirm infections the following primers were used: *Sy. kochii*, Skochii_F (GAGCTATACTCAAGTAAGTAATTG) and Skochii_R (CTTAATTTAGAGATATAGGTGTTTTC), annealing temperature of 50°C; *Sy. buchneri*, Sbuchneri_F (TTGAATTTTGGATGGATAATTTT) and Sbuchneri_R (CTCCATATAGTTAAAATCTTAC), annealing temperature of 50°C; *Sy. sp. ‘novel’*, Sunknown_F (GTATATTTAACTTTACTTCTCTTCAG) and Sunknown_R (CTGATTTATAACTGAACGTATTTC), annealing temperature of 45°C. All PCR reactions were run on a 1% agarose gel at 100V for 30-40 minutes to check for bands. Specimens that had more than 1% of amplicon reads assigned to *Symbiotaphrina* were screened with all three primer pairs. Some specimens could not be screened because all DNA was used during amplicon or UCE library preparation (specimens noted in Figure 3 as “nd”).

## Acknowledgements

We would like to thank Dr. Melissa Sanchez at the UNM Molecular Biology Core (MBC) Facility for quality control and sequencing services, and the UNM Center for Advanced Research Computing (CARC) for high performance computing resources used in this work.

## Data Availability Statement

The unprocessed sequencing datasets presented here can be found in the Sequence Read Archive (SRA) run by the US National Center for Biotechnology Information. The unprocessed UCE data is under SRA PRJNA1314812, and the unprocessed ITS amplicon data is under SRA PRJNA1317542. Processed data, such as the final ASVs, can be found at our Figshare at https://doi.org/10.6084/m9.figshare.33067949.

## Author contributions

AH and VGM designed the research. AH, VGM, TKP, TE, and RP collected, identified, and preserved insect specimens used in this manuscript. AH generated and analyzed the fungal ITS amplicon data and insect UCE data. AH and VGM wrote the first draft of the manuscript. All authors contributed feedback on the final manuscript draft.

