## Supplemental Figures for "Diffuse fungal symbiosis in Deathwatch beetles"

**Supplemental Figure 1.** Alternative phylogenies of beetle family Ptinidae using 95p dataset. Comparison of three different methods: a maximum-likelihood tree created with IQ-TREE2, a species tree from site patterns with SVDQuartets, and a species tree from gene trees with ASTRAL. All trees are based on Coleoptera-specific UCE loci recovered from at least 95 percent of specimens (95p matrix). To better denote differences between methods, the subfamilies are color-coded. Support values for IQ-TREE2 are bootstraps, for SVDQuartets are bootstraps, and ASTRAL are concordance vectors.

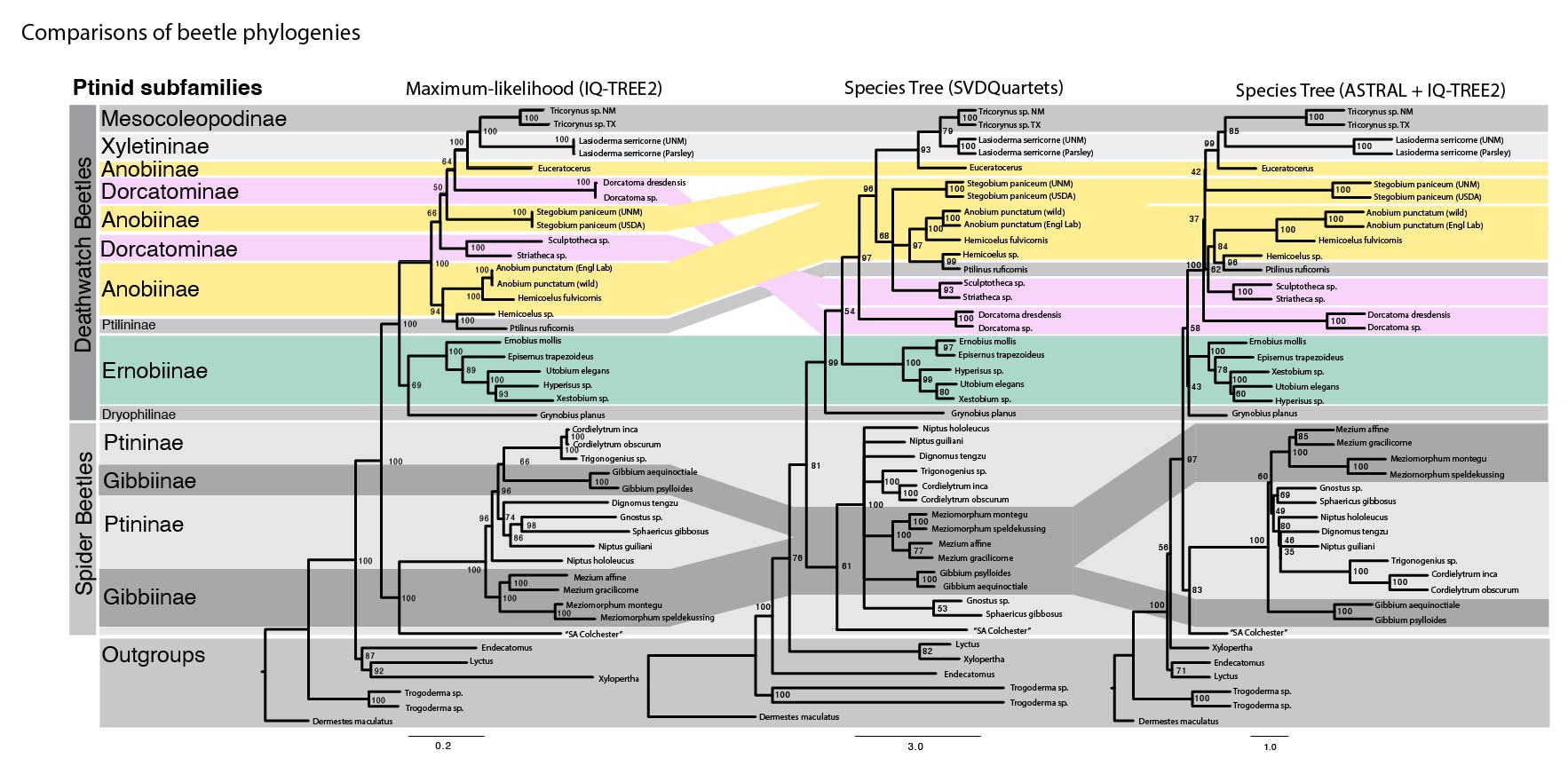

**Supplemental Figure 2.** Mirrored comparison of Maximum-likelihood trees generated with IQ-TREE2 using the 50p and 75p datasets. Taxonomic subfamilies are color-coded. Numbers represent bootstrap support values.

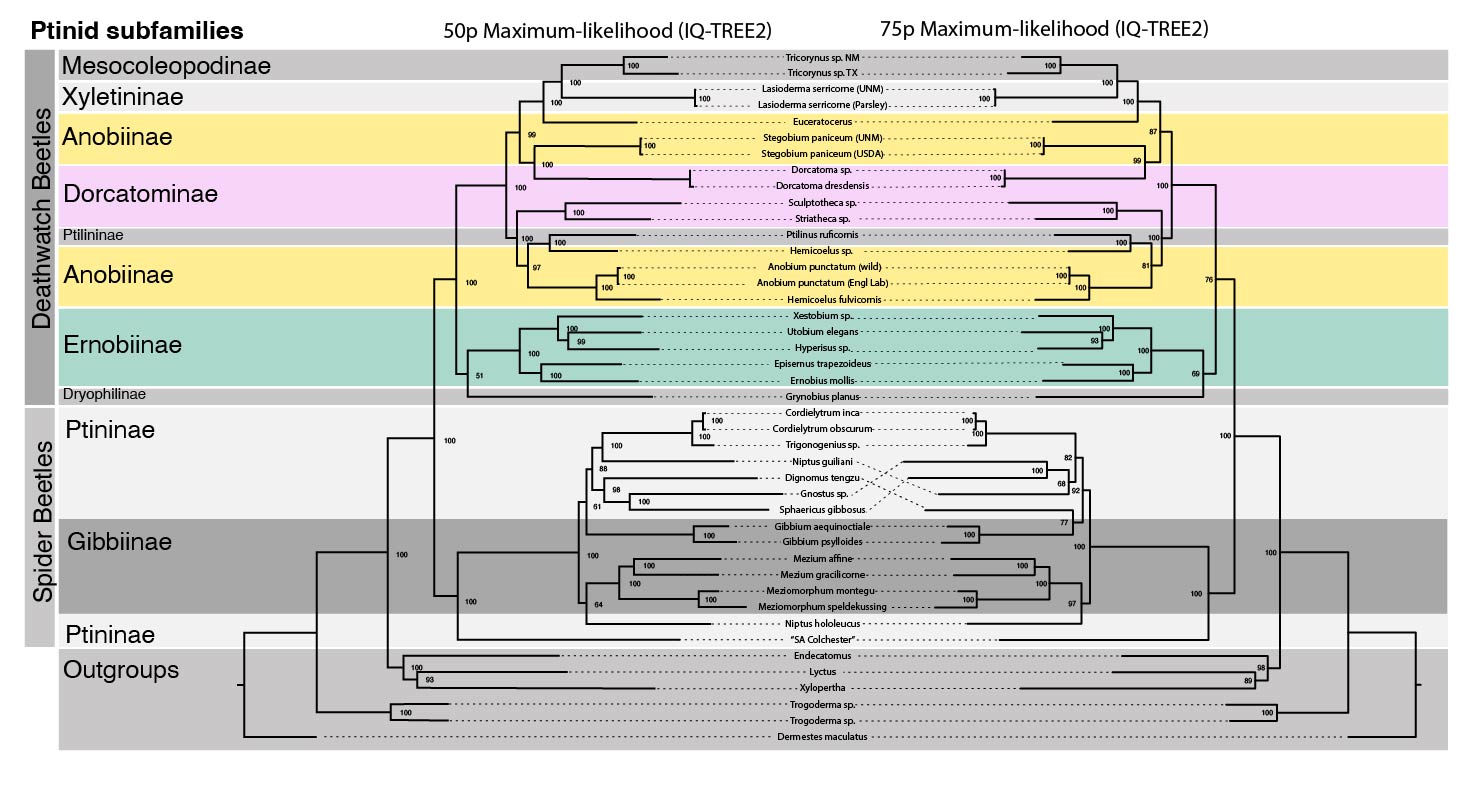

**Supplemental Figure 3.** Mirrored comparison of species trees generated with SVDQuartets using the 50p and 75p datasets. Taxonomic subfamilies are color-coded. Numbers represent bootstrap support values.

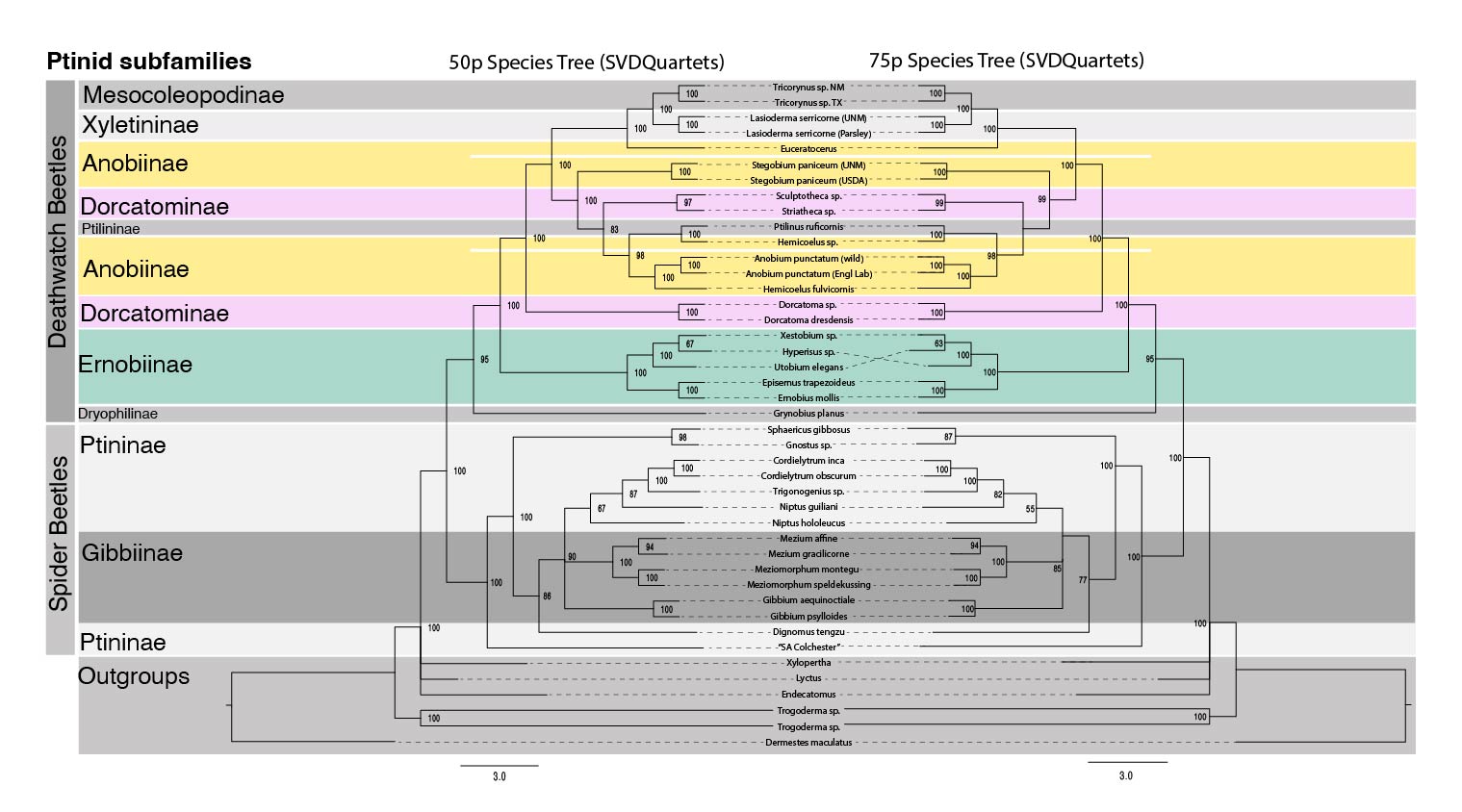

**Supplemental Figure 4.** Mirrored comparison of species trees generated with ASTRAL using the 50p and 75p datasets. Taxonomic subfamilies are color-coded. Numbers represent concordance vectors on a scale of 0-100 for better comparison to other methods presented in Figure 1 and S1-S3. Some dotted lines are emphasized to improve readability.
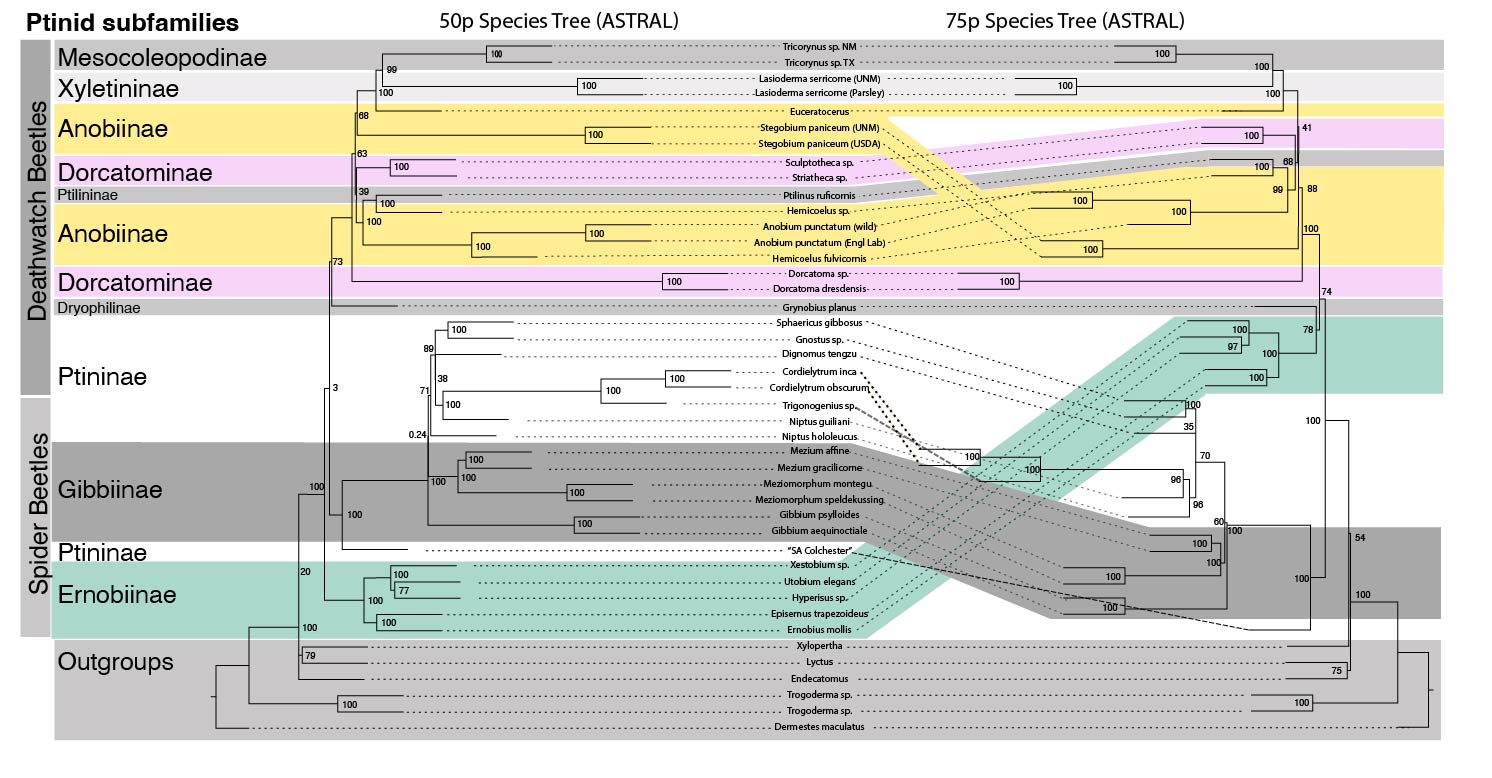

**Supplemental Figure 5.** Phylogenetic trees for *Nakazawaea* (A) and *Meyerozyma* (B) using trimmed ITS sequences. Sequences from this study are highlighted in gray. Support values are bootstraps.

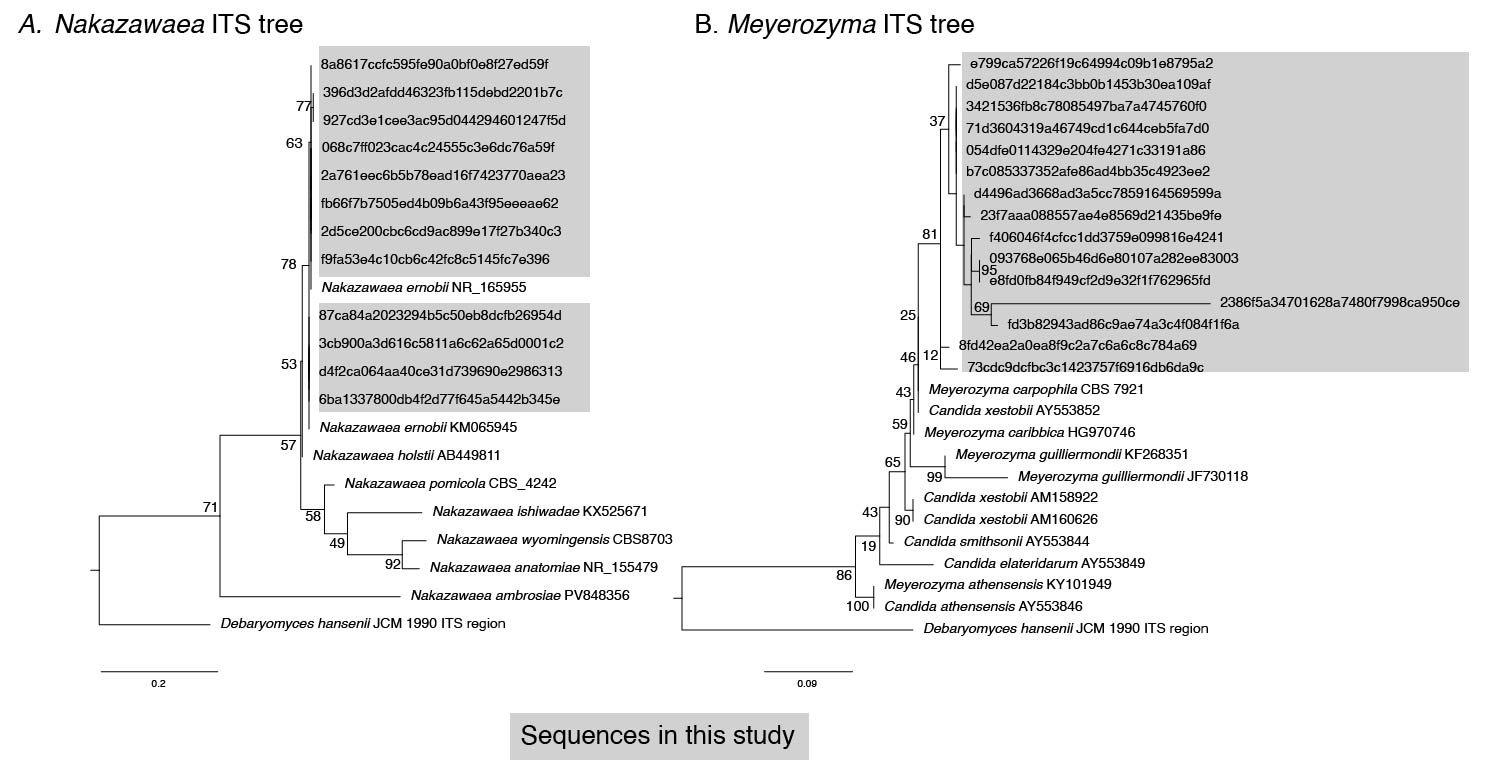

**Supplemental Figure 6:** Images of 1.5% agarose gels for species-specific *Symbiotaphrina* PCR screens. (A) Test of primers against DNA extracted from pure fungal cultures. (B) Screens of beetle DNA. Each column represents one DNA sample from one individual beetle specimen. Each row is a different primer set, specific to a *Symbiotaphrina* species.
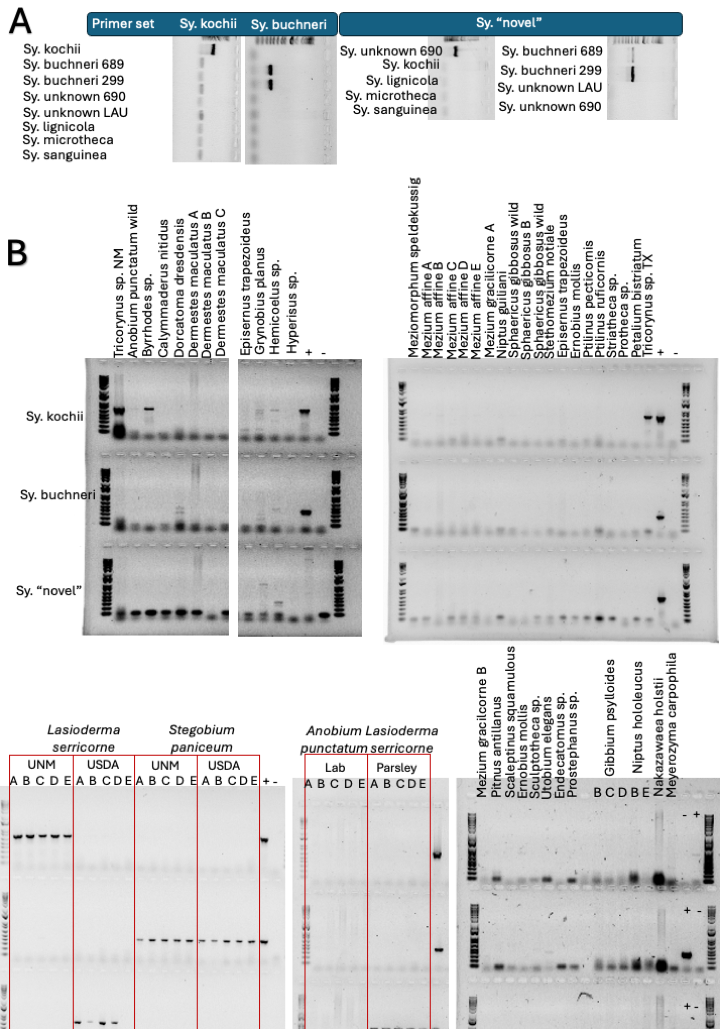

**Supplemental Figure 7.** Phylogenetic tree for the unidentified Helotiales spp. using trimmed ITS sequences. Sequences from this study are highlighted in gray. Support values are bootstraps.

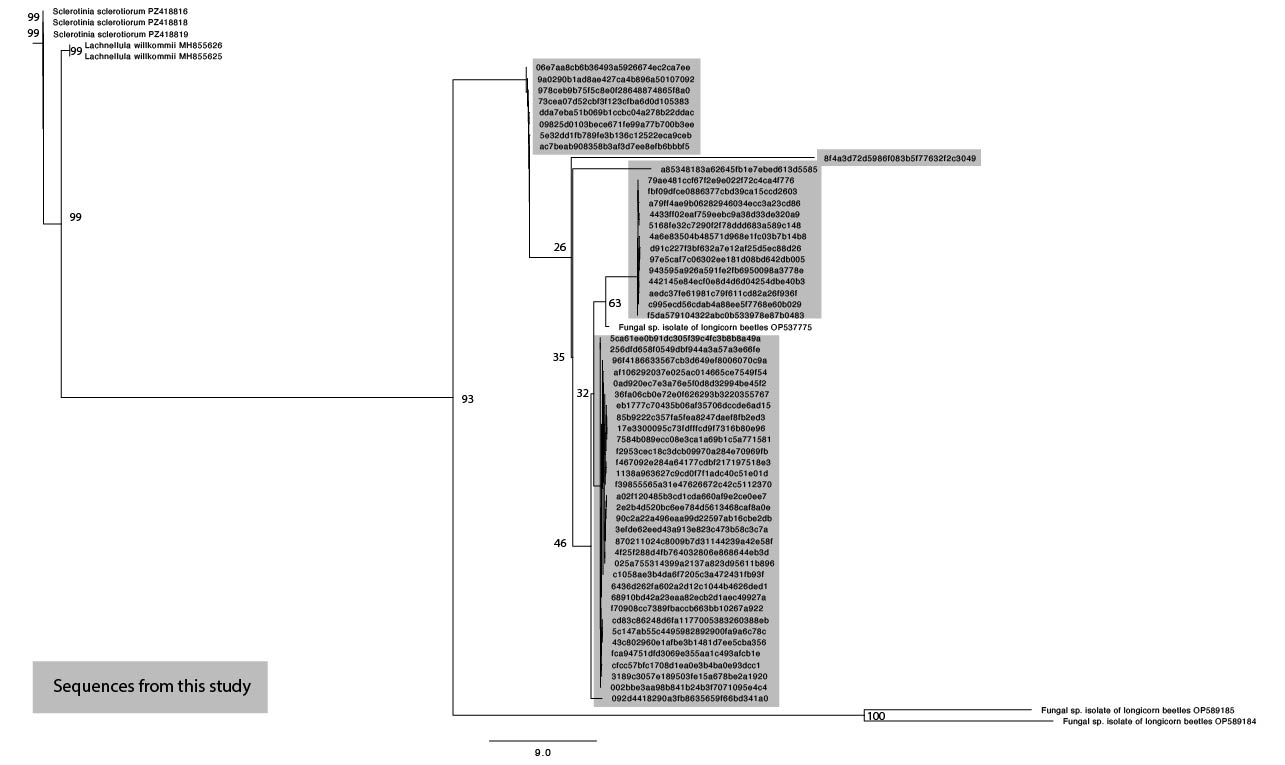

**Supplemental Figure 8:** Relative abundance of ITS ASVs grouped by taxonomic Family. Species highlighted in Figure 4 are grouped together here and colored gray.
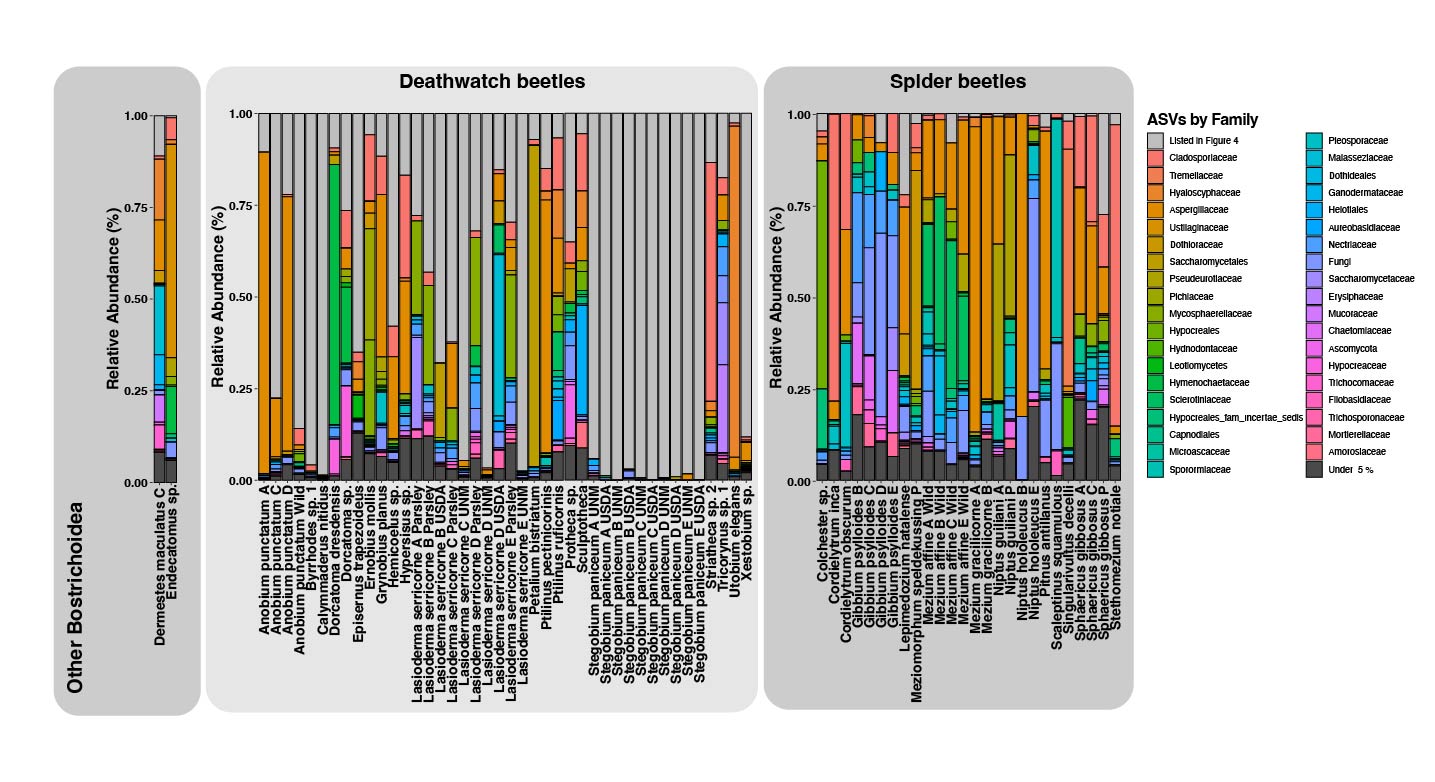
